# PepCentric Enables Fast Repository-Scale Proteogenomics Searches

**DOI:** 10.1101/2025.02.24.639867

**Authors:** Fengchao Yu, Andy T. Kong, Yi Hsiao, Alexey I. Nesvizhskii

## Abstract

Identifying novel peptides arising from alternative splicing, mutations, or non-canonical translations is a crucial yet challenging aspect of proteogenomics. We introduce PepCentric, a scalable computational platform and a web-based portal utilizing advanced 2-D fragment indexing for rapid peptide-centric searches across extensive mass spectrometry datasets. With robust false discovery rate control and optimized search performance, PepCentric offers an efficient tool for validating novel peptides and exploring proteomic variations. In a matter of seconds, users can search their novel peptides or proteins against 2.3 billion spectra collected from 66700 mass spectrometry runs, making it practical to rapidly validate proteogenomic hypotheses.

---

Proteogenomics is an emerging interdisciplinary field that integrates proteomics and genomics to provide a comprehensive understanding of cellular processes at the molecular level. A critical aspect of proteogenomic research involves the identification of novel peptides absent from reference protein databases^1,2^ using mass spectrometry (MS) data. These peptides can originate from alternative splicing^3^, gene mutations^4^, or non-canonical translation events^5^, each contributing to proteome complexity and functional diversity. Characterization of the “dark proteome” is important for advancing our understanding of complex diseases, including cancer^6^.

To detect novel peptides a commonly used approach^1,2,7^ involves building a custom protein sequence database (e.g. from genomics data acquired on the same samples) and searching acquired tandem mass (MS/MS) spectra against it using traditional peptide identification tools. However, this approach is inefficient when handling large datasets, as it requires downloading all files, generating custom databases, and conducting extensive searches, which makes it both time-consuming and error prone. Additionally, controlling the false discovery rate (FDR) globally becomes challenging when datasets are searched separately. Furthermore, a biologist studying a particular disease may be interested in searching for evidence of the expression of a specific candidate peptide or protein, without needing to reprocess any MS data from scratch to perform a new search. To address these limitations, researchers have attempted to build MS databases and query peptides against them^8-10^, with recent studies showcasing various applications of peptide-centric proteogenomics searches^8^. However, these earlier attempts either required specialized hardware^10^ or, due to the lack of optimized indexing techniques^8,9^, suffered from slow processing speeds and limited scalability.

Here, we present a computational solution that enables querying any novel peptide(s) of interest against a vast repository of MS data in milliseconds, making it as fast and easy as performing a Google search. We describe a method for indexing repository-scale MS spectra and an associated web-based infrastructure that allows users to input proteins or peptides and search against indexed data. Our approach is rooted in our previous discovery of a fragment ion indexing algorithm^11^, and the search engine MSFragger that dramatically changed the speed of conventional peptide identification by indexing theoretical fragmentation spectra prior to the search^11,12^. To enable peptide-centric searches, we implement a reversed strategy of indexing the experimental MS data. To overcome the technical hurdles of dealing with large indexes that cannot fit into conventional memory, we developed a 2-D fragment indexing technique that is compatible with commodity hardware while maintaining fast speed. Here, we describe our computational infrastructure, named PepCentric, and make a large collection of indexed MS data available for proteogenomics queries on our public web serve http://peptidecentric.org:8080/

PepCentric is designed for fast searching of repository-scale MS data and retrieval of associated metadata by indexing all levels of information, including run names, spectrum identifiers, spectrum precursor masses, fragment m/z values, and fragment intensities. The current version of the PepCentric webserver contains over 200 billion peaks from more than 2.3 billion MS/MS spectra collected from 66700 mass spectrometry runs downloaded from public data repositories including PRIDE^13^, MassIVE (https://massive.ucsd.edu/), Proteomic Data Commons^14^ (PDC), and BioPlex^15^. To efficiently search peptides against these data, we designed a novel 2-D fragment indexing approach that utilizes a tailored filesystem-less data layout and a customized caching system (**Figure 1a**). This approach involves converting the fragment m/z value into a fixed-point unsigned 32-bit number and dividing it into two parts: higher 16 bits and lower 16 bits. The higher part is used as the index of the M/Z bucket, which contains fragment peak entries that include a global spectrum identifier, lower portion of the fragment peak M/Z, and normalized peak intensity. When writing the fragment index to hard drives, the fragment peaks in each bucket are assigned to blocks whose size equals the smallest input/output unit of a solid-state drive (SSD). After writing all the blocks and buckets to the SSD, the bucket offset is used as the first-dimensional index, while the block identifier represents the second-dimensional index. Most operating systems and filesystems include an integrated caching and prefetching system to aid data retrieval. This can improve the speed of sequential data access, but when used to retrieve random data, such as what is extensively done in PepCentric searching, it leads to inefficiencies and increased latency. To address this issue, we bypassed these mechanisms by laying out the data structure of our 2-D fragment index directly on the disk. During the spectral searching process, hyperscore, expectation value (e-values), and mass difference are calculated for each peptide-spectrum matches (PSM) as previously described.^11,12^

**Figure 1.**
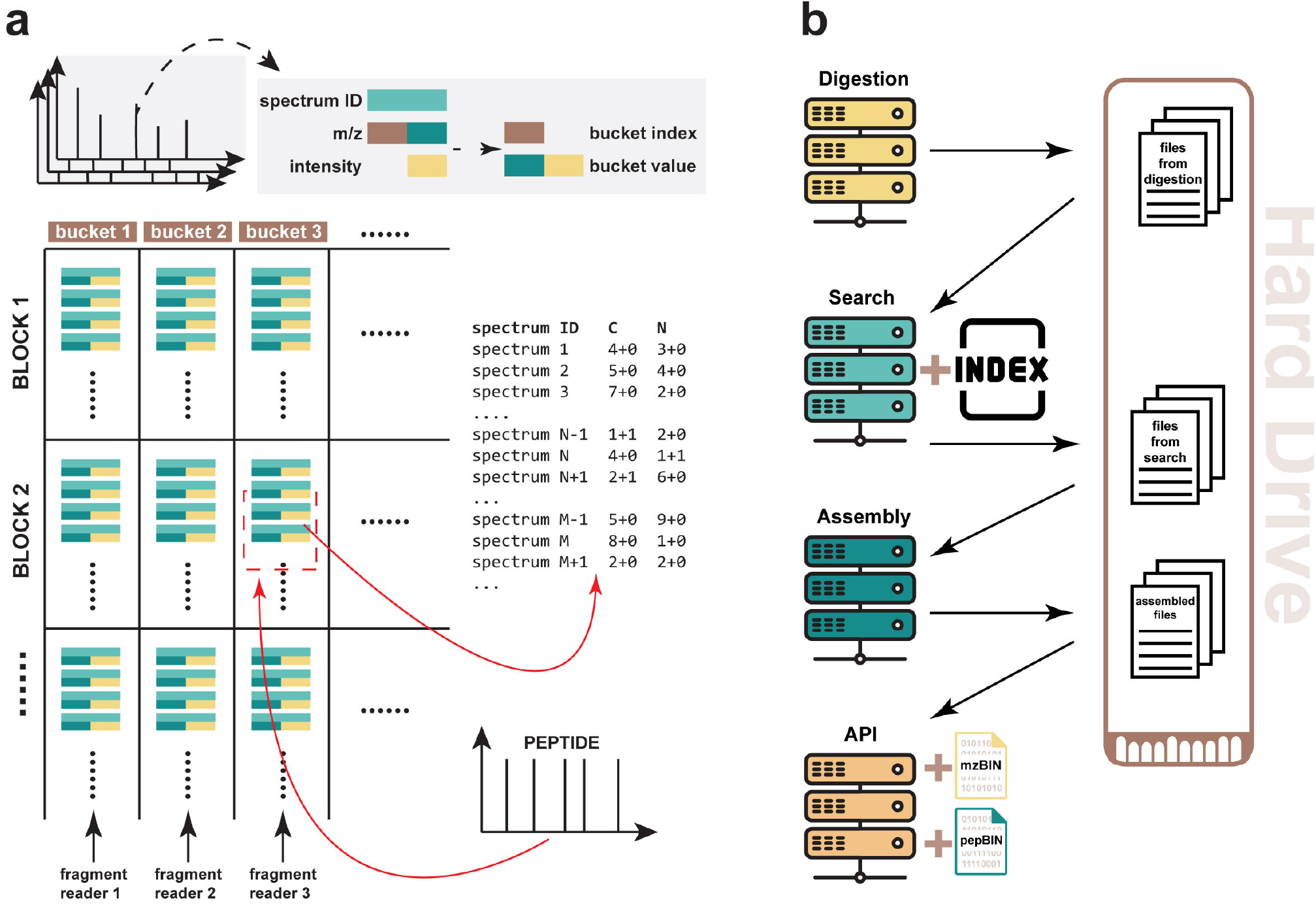
2-D fragment indexing and backend server architecture. **(a)** Illustrations of **(a)** Schematic representation of the 2-D fragment indexing method. The upper part illustrates the conversion of experimental fragment peaks into numerical arrays optimized for 2-D indexing. The lower part is a diagram of the 2-D index structure and the query process (red arrows) for a given peptide. **(b)** Diagram of the backend server architecture, including digestion, search, assembly, and API servers. These servers communicate via a hard drive. The search server accesses index data stored on a high-speed hard drive, while the API server retrieves mzBIN and pepBIN files to provide data for result presentation.

To maintain the quality and reliability of reported peptide matches, it is essential to control the false discovery rate (FDR). While the hyperscore measures the similarity between the experimental and theoretical peptide fragmentation spectrum, neither the hyperscore nor the e-value (which is a normalized hyperscore) provide any global-level statistical metrics^16^. Thus, we developed a PepCentric module to calculate global p-value at the PSM, peptidoform (sequence + modifications), and peptide (i.e. unique sequence) levels, and also FDR (q-value) at the peptide level. We conducted a search using over 4 million decoy peptides against the fragment index to generate a null distribution of e-values (which is used as the primary score for each PSM), saved in a binary file. Then, for each peptide sequence queried by the user using PepCentric, a null distribution is retrieved and used to calculate the p-value. The Benjamini-Hochberg procedure is used to calculate the FDR, and transform it into the q-value, providing a very conservative measure when querying multiple peptides simultaneously (see **Methods**).

To make PepCentric widely accessible, we designed a server with two main components: frontend and backend. The frontend comprises a website and a structured query language (SQL) database that records user queries (**Figure 2a**). The backend comprises several servers, including a digestion server (the interface allows querying protein sequences that are internally digested into peptides), a search server, an assembly server, and an API server (**Figure 1b**). The digestion server interacts with the SQL database to retrieve user-provided sequences and performs *in silico* digestion. The search server then searches the digested sequences against the indexes. The assembly server collects and summarizes the search results, while the API server communicates with the frontend to provide the search results, annotated spectra, and other relevant information. The infrastructure supports querying multiple separate indexes, and can engage multiple search servers. The index files are stored on a dedicated SSD to enable fast data retrieval. Furthermore, PepCentric also stores peptide sequences and identification scores obtained by MSFragger closed and open searches of the MS/MS spectra against the reference proteome database (UniProt, reviewed sequences). MSFragger searches are performed at the index preparation step, and the results are stored and then retrieved by PepCentric to be shown alongside the matches obtained for the user’s query. This enables the user to perform manual inspection to determine, for each candidate novel PSM, how much the novel peptide match is better than the best reference PSM. The availability of open search results further helps to identify possible misidentifications of novel peptides resulting from common mass shifts (e.g., chemical or biological modifications) that were unaccounted for in the search - a common but underappreciated issue in proteogenomics searches^1^.

**Figure 2.**
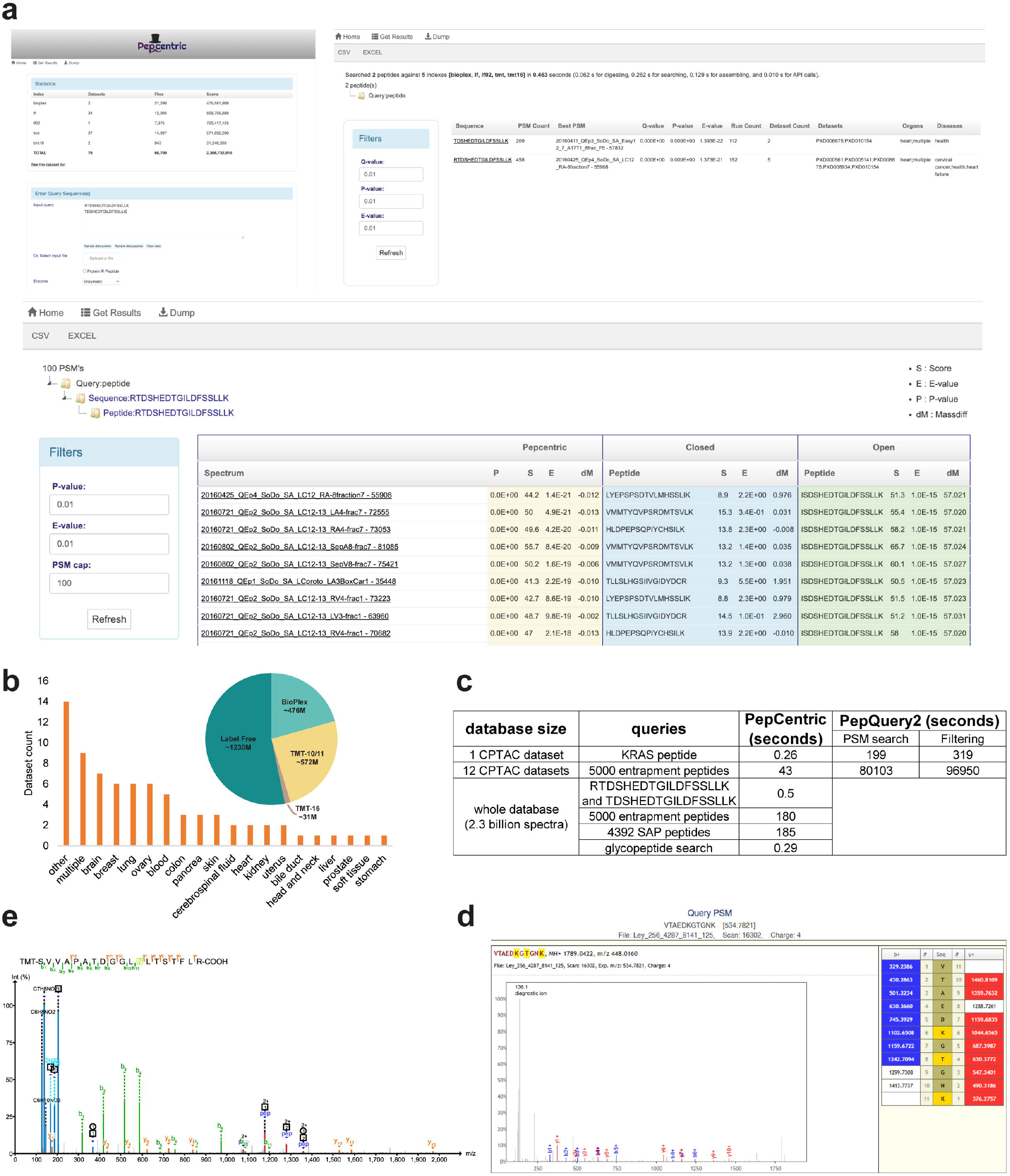
Frontend webpage illustrations and performance evaluations. **(a)** Illustrations of the webpages interface, including a landing page for user input and two result pages displaying reports at the peptide sequence level and peptidoform level, respectively. **(b)** Summary of the datasets in PepCentric. **(c)** Runtime comparison using various queries and datasets. PepQuery2’s runtime is divided into “PSM search” (searching the query against the spectra) and “Filtering” (refining PSMs using reference databases and modifications from the UniMod database). **(d)** A screenshot of the visualization page displaying the identified AMPylated peptide VTAEDKGT[329.052520]GNK. Shown matched fragments account for the AMPylation-specific neutral losses. **(e)** An MS/MS spectrum of the identified glycopeptide SVVAPATDGGLN[1647.6085]LTSTFLR. The spectrum was extracted from the PepCentric database and visualized using PDV.

At present, PepCentric contains four sets of indexes: TMT-10/11, TMT-16, BioPlex, and Label free (**Figure 2b, Supplementary Data 1**). Collectively, there are ∼2.3 billion MS/MS spectra from 28 disease types and healthy samples (**Supplementary Data 2**), 19 human tissues and cell lines (**Figure 2b, Supplementary Data 3**). We plan to continually add new datasets and maintain the server in the long run. In addition to the web server, we have developed a RESTful API to enable programmatic access.

To evaluate FDR control in PepCentric, we generated 5000 entrapment peptides by random shuffling of amino acids in peptide sequences obtained by *in silico* digestion of the *Homo sapiens* protein sequence database (**Methods**) and searched them using PepCentric. When searching against 12 large CPTAC datasets, PepCentric reported 88 peptides passing the 0.01 global p-value cutoff, however, none passed the stringent 0.01 global q-value filter (**Supplementary Data 4**). In comparison, PepQuery2, when searching same peptides against the same 12 datasets, reported 893 entrapment peptides as “confident” using the same test (**Supplementary Data 5**), further highlighting the need for FDR control. We also compared the runtimes between PepCentric and PepQuery2 using an Amazon Elastic Compute instance (i4g.4xlarge, AWS Graviton2 CPU, 16 cores, 128 GB memory) and observed a ∼4000 times faster overall analysis time of PepCentric compared to PepQuery2 (∼2000 times faster if only considering PepQuery2’s PSM searching step) (**Figure 2c**). When we searched the same 5000 entrapment peptides against the entire collection of ∼2.3 billion spectra, PepCentric still reported 99 entrapment peptides with 1% global p-value, and 0 with 1% global q-value (**Supplementary Data 6**). This reflects the highly conservative nature of the Benjamini-Hochberg adjustment, and suggests that, in certain applications, a more relaxed q-value filter of e.g. 0.05 may be applicable. Notably, PepQuery2 could not search 5000 peptides against the entire dataset in a reasonable time. Overall, these experiments show robust FDR control and excellent computation speed of PepCentric, enabling practical and routine interrogation of novel peptides and proteins at a massive, “repository-wide” scale.

After demonstrating that PepCentric has low FDR, we evaluated its sensitivity by replicating the KRAS G12D mutated peptide search experiment performed by Wen et al^8^ (**Methods**). We searched the KRAS G12D peptide (LVVVGA**D**GVGK) against the same 12 cancer datasets used by Wen et al^8^. Using a combination of 1% global q-value, PepCentric identified 30 PSMs, of which 28 overlapped with PepQuery2 (**Supplementary Data 7**). We manually inspected the two PSMs uniquely reported by PepCentric and found both proteome- and genome-level evidence. **Supplementary Figure 1** shows that the KRAS G12D peptide has a better match than the wild-type peptide as reported by PepQuery2. The genomics data also shows that the tumor sample C3N-00516_T, where one of those two PSMs were found, has (34-36)gGt>gAt genomic mutation corresponding to the G12D proteome mutation (**Supplementary Data 8**). Finally, the runtime comparison performed on the same computer showed that PepCentric was ∼2000 times faster than PepQuery2 in this experiment (∼800 times faster if only considering PepQuery2’s PSM searching step) (**Figure 2c**).

We next showcase how PepCentric can provide additional validation of novel peptides found in other studies. Lau et al^17^ highlighted the detection of a skipped exon (SE) splice junction peptide RTDSHEDTGILDFSSLLK in their human heart study. We searched this sequence using PepCentric (which does not contain the data generated as part of the Lau et al^17^ study) and detected it multiple human heart samples (**Supplementary Data 9**). We further applied PepCentric to a large-scale single-amino acid polymorphism (SAP) study^18^. The original publication reported 4392 tryptic peptides with SAP mutations. The PepCentric search reported 3558 peptides detected in various public datasets, which again did not include the original data (**Figure 2c, Supplementary Data 10**). Of note, PepCentric took only ∼3 minutes to search 4392 peptides against ∼2.3 billion spectra on a computer with a 16-core CPU and 128 GB memory. These analyses further demonstrate that PepCentric can be used as a valuable resource to validate other researchers’ novel peptide discoveries using complementary, publicly available MS datasets assembled as part of PepCentric.

Finally, we demonstrate that PepCentric can also be used to search peptides with post-translational modifications. In the first example, we searched for evidence of an AMPylated peptide VTAEDKGT[329.052520]GNK from Endoplasmic reticulum chaperone BiP (site T518). This rare modification is known to occur on several proteins, including BiP, and has been the subject of several in-depth studies^19^. PepCentric search has been able to confidently detect this AMPylation event in data from 583 runs from 31 studies (**Supplementary Data 11**). Manual inspection of the selected identified MS/MS spectra using PepCentric’s built-in spectral viewer further supported the validity of the identification through clear observation of diagnostic ions of AMPylation such as a peak at m/z 136.1 as well as signature losses of -347 Da (**Figure2d)**. Second, we sought to determine the profile of expression of a tryptic peptide SVVAPATDGGLNLTSTFLR from prostaglandin-H2 D-isomerase (PTGDS) containing a known N glycosylation site. Recent studies have highlighted the heterogeneity and role of N glycans on PTGDS in health and disease^20^. We searched SVVAPATDGGLN[1647.6085]LTSTFLR (the mass shift corresponding to a N-glycan H4N5F2S0) using PepCentric. This glycoform was confidently detected in 87 runs from 8 datasets, which were predominantly, as expected, cerebrospinal fluid (CSF) and brain tissue samples (**Supplementary Data 12**). In addition to the built-in viewer, PepCentric can export spectra to be annotated and visualized using PDV^21^. **Figure 2e** illustrates this functionality using one of the PDV-annotated MS/MS spectra, which shows good fragment ion coverage of the peptide, as well as oxonium ions and Y-ions expected in the glycopeptide spectra.

In conclusion, PepCentric represents a significant advancement in the field of proteogenomics by enabling fast, scalable, and accurate peptide-centric searches across extensive mass spectrometry datasets. By leveraging advanced 2-D fragment indexing, PepCentric overcomes the previous limitations of speed and FDR control, providing researchers with a powerful tool for validating novel peptides and exploring proteomic variations. Its web-based platform ensures accessibility and facilitates the validation of proteogenomic discoveries as well as the search for rare PTMs, making it an invaluable resource for advancing our understanding of complex diseases and cellular processes.

## METHODS

### All levels indexing enables fast searching in repository-scale

Although PepCentric builds upon the fragment indexing strategy pioneered by MSFragger, the algorithms are reengineered to address the challenges of repository-wide peptide-centric searching. First, it indexes several levels of data, including LC-MS files, spectra metadata, peptide identification scores, precursor ions, and fragment peaks. When preparing the indexes, MSFragger searches the spectra (closed and open search) against the reference proteome database to obtain canonical peptide matches. These scores are stored in pepXML format and used as references after PepCentric searches for novel peptides. MSFragger also calibrates the masses and writes the spectra to mzBIN files. The mzBIN is an indexed binary format that is fast for random access. Second, the fragment index is stored on hard drives to eliminate size constraints from the RAM. However, solid state drives have slower input/output (IO) speed compared to RAM. PepCentric implements a new fragment indexing strategy to reduce the IO and maintain the speed, coupled with a new search algorithm leveraging the revised indexing and parallelization to push the searching speed to the hard drive’s limit.

Given a dataset of MS spectra, PepCentric takes the mzBIN and pepXML files (from the initial MSFragger search, as described above) as inputs to extract information and build the indexes. PepCentric first converts the pepXML files to a pepBIN binary format that has the PSMs indexed and encoded to make random access faster compared to XML formatted files. The pepBIN files will also provide information about matches against the reference database for each spectrum that corresponds to novel peptides during the result presentation stage. Since the number of spectra matching the novel peptides can be substantial, the indexed and binary pepBIN format significantly accelerates this process. Then, PepCentric extracts run names (i.e., the file name of each LC-MS injection), spectrum names, spectrum precursor masses, and spectrum precursor charges from the mzBIN files. These values are sorted based on their precursor masses and stored in a binary form. PepCentric does not use SQL database because it has larger file size and slower querying speed compared to our binary formats optimized for our specific purposes. From the pepBIN files, PepCentric extracts all spectra’s canonical (i.e., in reviewed UniProt) peptide matches, hyperscores, e-values, and delta masses that are the mass differences between the precursor mass and calculated peptide mass. For the MSFragger open search output, PepCentric puts the delta mass, which could correspond to a modification, on each amino acid to calculate the localization-aware hyperscores^12^. These scores are indexed and stored in binary form. Note that the MSFragger search results are only used to provide the user with additional information regarding reference peptide assignments as part of the PepCentric report; they are not used as part of the PepCentric search itself.

After indexing the run- and spectrum-level information, PepCentric extracts and indexes the fragment peaks of all MS/MS spectra. There are hundreds of billions of peaks to be written to the hard drives during the index building stage, and the program locates and retrieve the peaks from the hard drives during the PepCentric searching stage. Compared to the fast central processing unit (CPU) speed, the IO between the hard drive, RAM, and CPU is the bottleneck. To minimize the IO latency, we developed a 2-D indexing method coupled with a customized filesystem and caching system (**Figure 1a**). Given a fragment M/Z, PepCentric converts it into a fixed-point unsigned 32-bit number. Then, it splits its higher and lower 16 bits into two parts. The higher part is used as the index of the M/Z bucket (first-dimensional index). In each bucket, there are fragment peaks with closed M/Z from billions of spectra. Within each bucket, we developed another index to quickly locate the peaks within a given precursor mass range. Each fragment peak has three fields: the global spectrum identifier, the lower 16 bits of the peak M/Z, and the normalized peak intensity encoded in 16 bits. Because the global spectrum identifier is sorted based on the precursor mass, the fragment peaks are sorted based on their global spectrum identifier. When writing the 2-D index to dedicated hard drives, PepCentric writes each bucket separately. In each bucket, the fragment peaks are assigned to blocks with a size of 4096 bytes, which is the smallest unit of IO for most solid-state drive (SSD). The first spectrum identifier is used as the block identifier. After writing the first bucket, PepCentric records the bucket offset, writes the second bucket, and keep going until writing all buckets. The bucket offset corresponds to the first-dimensional M/Z bucket index, and the block identifier is the second-dimensional index.

In practice, most operating systems (OS) have built-in caches and prefetching to improve IO performance. When reading data sequentially, such practices can improve speeds. However, when reading random data like what PepCentric searching does, it only increases overhead costs and latencies. Thus, we bypassed all such mechanisms by directly writing the fragment index to hard drives without the use of OS’ existing filesystems. Given a fragment M/Z, PepCentric locates the starting blocks and iteratively reads only the necessary blocks for peak matching. Together with the 2-D indexing, PepCentric can search billions of MS/MS spectra in seconds.

Hyperscore, e-value, and delta mass are calculated to measure the similarity of the peptide-spectrum matches (PSM) during the PepCentric search. A peptide’s hyperscore is the highest value among its PSMs. A peptide’s e-value is the lowest (best) value among all corresponding PSMs. The peptide sequence for a modified peptide is obtained after stripping the modifications. Then, the reported hyperscore and e-value score are the best ones among the peptides with the same sequence.

### Global false discovery rate control in PepCentric

False discovery rate (FDR) control is critical for the quality and confidence of the reports. Almost all queries have matches from the PepCentric search. The hyperscore and e-value are measures of the similarity between the peptide and the matched spectrum. However, they do not reflect global statistical confidence. We developed modules to calculate global p-values, FDRs, and q-values at the peptide, sequence, and PSM levels. We first generate a null distribution by searching a large number (millions) of non-existing peptides against the fragment indexes. After searching, the peptide sequence e-values are extracted and stored in a binary file. When searching a new peptide against the fragment indexes, the p-value is calculated as follows:

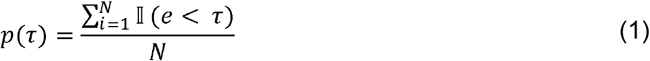

where N is the number of e-values in the null distribution, 𝕀 is the indicator function, *e* is the e-value in the null distribution, and *r* is the e-value of a given match. Given the p-value, we can then calculate the sequence-level FDR using a method similar to Benjamini-Hochberg procedure^22^

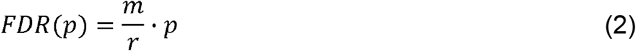

where *m* is the total number of querying sequences, *r* is the rank of a given p-value, and p is the p-value. Finally, we convert the FDR to q-value^23^

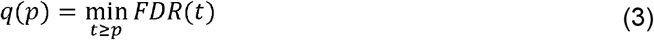

### Downloading, searching, and indexing the large-scale projects from the public repositories

We have downloaded and indexed in total 66 projects from PRIDE, MassIVE, Proteomic Data Commons (PDC), and BioPlex repositories. There are in total 66700 LC-MS files, and ∼2.3 billion tandem mass spectra. They were first searched by MSFragger using closed and open searches. The FASTA database contains human proteome from UniProt (https://www.uniprot.org/proteomes/UP000005640) and 116 common contaminants from cRAP (https://www.thegpm.org/crap/). For label-free datasets, protein N-terminal acetylation and oxidized methionine were set as variable modifications. Carbamidomethyl on cysteine was set as a fixed modification for non-BioPlex datasets. For the TMT datasets, protein N-terminal acetylation, oxidized methionine, and peptide N-terminal TMT were set as variable modifications. Carbamidomethyl on cysteine and TMT on lysine were set as fixed modifications. For closed and open searches, the maximum allowed missed cleavages were set to 2 and 1, respectively. Detailed MSFragger parameter files can be found in **Data availability**. The spectra and the search results were indexed and deployed on the PepCentric server.

### RESTful API to perform PepCentric search programmatically

In addition to the straightforward website as the user interface, we developed RESTful APIs to allow people to use PepCentric programmatically. With the APIs, people can submit peptides and proteins and retrieve results. A detailed documentation can be found in **Supplementary Notes**.

### Entrapment peptides search

To evaluate the accuracy of the FDR control, we generated 5000 nonexistent peptides and searched for them using PepCentric and PepQuery2, respectively. *Homo sapiens* proteins from UniProt (downloaded on November 27, 2020) and common contaminant proteins from cRAP (https://www.thegpm.org/crap/) were used to *in silico* generate tryptic peptides without any missed cleavages and requiring length of 7 amino acids or more. The starting set contained 626755 peptides from 75117 proteins. The peptides were randomly shuffled with C-terminus fixed. Shuffled peptides with sequences matching the proteins in the original database were discarded. Finally, 5000 shuffled peptides were randomly selected as entrapment peptides. Since searching 5000 peptides against the whole PepCentric dataset using PepQuery2 is infeasible, we selected 12 datasets from 10 Clinical Proteomic Tumor Analysis Consortium (CPTAC) cohorts used in the original PepQuery2 publication^8^. The peptides reported by PepQuery2 with a tag “confident” equals “yes” were considered as confidently identified. For PepCentric, we searched 5000 entrapment peptides against the same 12 datasets, as well as the whole database containing ∼2.3 billion spectra. The peptides were filtered as described in the main text.

### PepCentric searches

PepCentric was used to search the KRAS G12D mutated peptide, LVVVGA**D**GVGK, against 10 CPTAC cohorts from 12 datasets. The same mutated peptide and datasets were also used by Wen et al^8^. We ran PepCentric and PepQuery2 on the same computer in Amazon Elastic Compute Could (i4g.4xlarge, AWS Graviton2 CPU, 16 cores, 128 GB memory), respectively. For PepQuery2, the PSMs with “confident” equals to “yes” were used as the result. For PepCentric, the PSMs were filtered with 1% global q-value. The skipped exon (SE) splice junction peptide, RTDSHEDTGILDFSSLLK, and its non-missed-cleavage version, TDSHEDTGILDFSSLLK, the glycopeptide SVVAPATDGGLN[1647.6085]LTSTFLR and the AMPylated peptide VTAEDKGT[329.052520]GNK were searched in the same way. To generate annotated MS/MS spectra, selected MS/MS scans were extracted from the original mzML files and visualized using PDV^21^. Single-amino acid polymorphism (SAP)-mutated peptides were downloaded from Sinitcyn et al^18^. In total 4392 non-redundant peptides were searched and filtered as described above.

## Data availability

The raw LC-MS data used in this study was downloaded from the public repository. The detailed list can be found in http://peptidecentric.org:8080/datasets. The proteome database file is available in the UniProt database under the proteome ID UP000005640 (*Homo sapiens*). The Genomics data for the KRAS G12D mutation is available in https://proteomic.datacommons.cancer.gov/pdc/cptac-pancancer (file name “Mutation_Broad_WashU_union_v1.zip”). **Source data** are provided with this paper.

## Acknowledgements

This work was funded in part by NIH grants R01-GM-094231, U24-CA210967, and U24-CA271037 (to A.I.N.). We thank Snehal Patil for technical assistance with the development of the web interface.

