## Supplementary Notes for "PepCentric Enables Fast Repository-Scale Proteogenomics Searches"

**RESTful API**

- <backend server url> is addresss:port. Port's default value is 50001.
- <backend server url>/submit?email=<email>&jobName=<job name>&jobType=<job type>&query=<query>&enzyme=<enzyme>. Note: email and jobName are optional. <job type> can be peptide and protein. query is the peptide or protein (including protein header) sequences. enzyme can be enzymatic and nonspecific.
- <backend server url>/jobstatus?jobID=<uuid>
- <backend server url>/result?jobID=<uuid>
- <backend server url>/showSequence?jobID=<uuid>&proteinID=<protein ID>&q=<q-value>&p=<p-value>&e=<e-value>
- <backend server url>/showPeptide?jobID=<uuid>&proteinID=<protein ID>&sequenceID=<sequence ID>&p=<p-value>&e=<e-value>
- <backend server url>/showSpectrum?jobID=<uuid>&proteinID=<protein ID>&sequenceID=<sequence ID>&peptideID=<peptide ID>&psmCap=<PSM cap>&p=<p-value>&e=<e-value>
- <backend server url>/countSpectrum?jobID=<uuid>&proteinID=<protein ID>&sequenceID=<sequence ID>&peptideID=<peptide ID>&p=<p-value>&e=<e-value>
- <backend server url>/searchresults?runID=<file name>&scanNum=<scan number>
- <backend server url>/dump?jobID=<uuid>&q=<q-value>&p=<p-value>&e=<e-value>
