## Supplementary Figures for "PepCentric Enables Fast Repository-Scale Proteogenomics Searches"

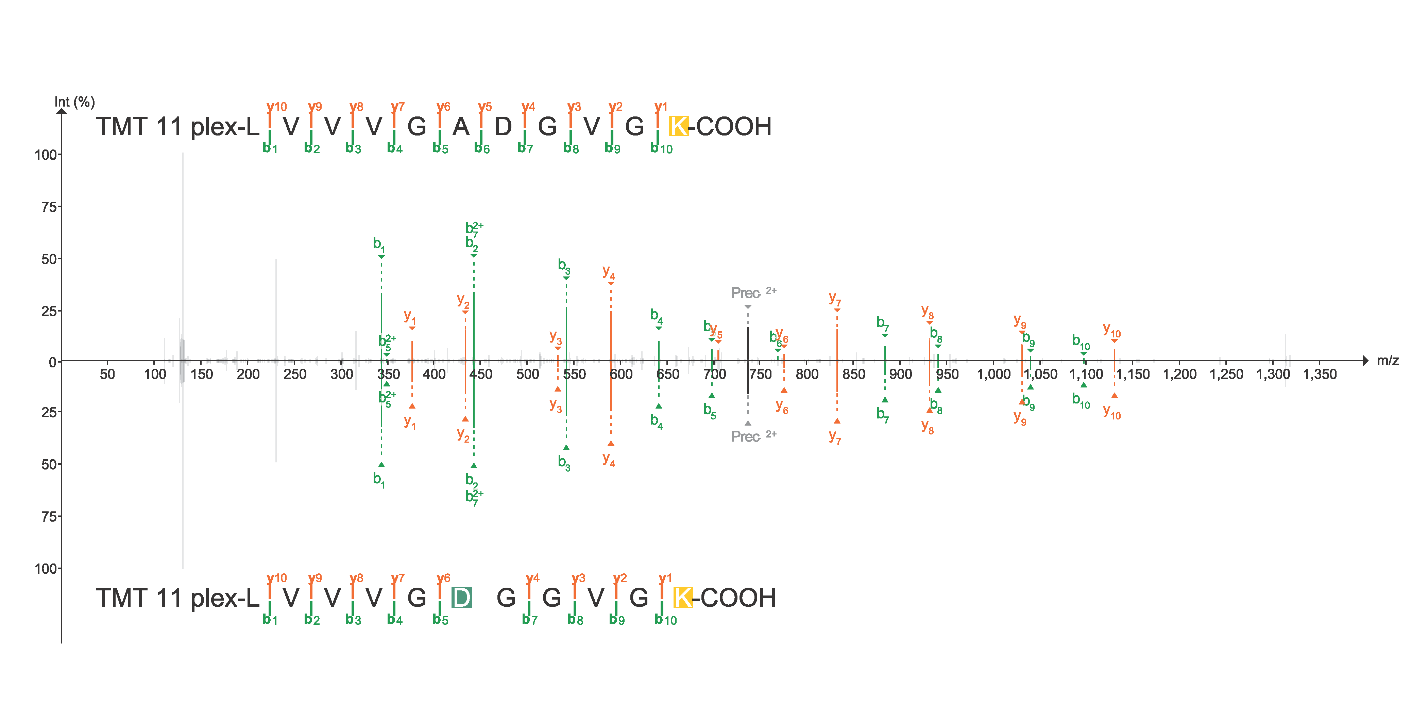

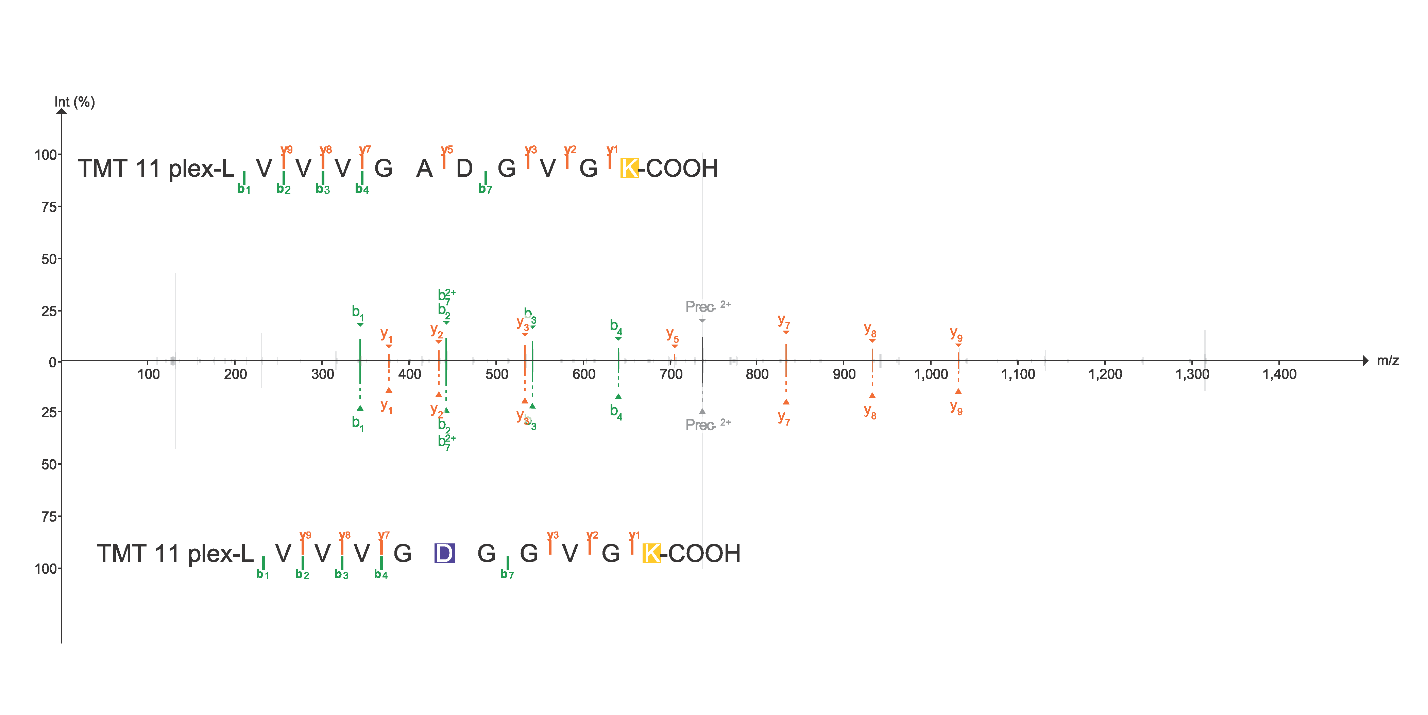

**Supplementary Figure 1.** Two annotated spectra showing the PSMs should not be discarded by PepQuery2. The upper panel is from scan “10CPTAC_PDA_W_JHU_20191207_LUMOS_f16.27993.27993.2” and the lower panel is from scan “16CPTAC_UCEC_W_PNNL_20180503_B4S4_f12.26903. 26903.2”. In both scans, PepQuery2 discarded the peptide “n(229)LVVVGADGVGK(229)” because another peptide “n(229)LVVVGD(14)GGVGK(229)” scored better. However, the annotated spectra show that there are two (upper panel) and one (lower panel) more fragments matching to “n(229)LVVVGADGVGK(229)”.
